# Genetic analysis of *Aedes aegypti* captured in two international airports serving to the Greater Tokyo Area during 2012—2015

**DOI:** 10.1101/823138

**Authors:** Kentaro Itokawa, Jinping Hu, Nayu Sukehiro, Yoshio Tsuda, Osamu Komagata, Shinji Kasai, Takashi Tomita, Noboru Minakawa, Kyoko Sawabe

## Abstract

Introduction of exotic diseases vectors into a new habitat can drastically change the local epidemiological situation. During 2012—2015, larvae and an adult of the yellow-fever mosquito, *Aedes aegypti*, were captured alive in two international airports serving to the Greater Tokyo Area, Japan. Because this species does not naturally distribute in this country, those mosquitoes were considered to be introduced from oversea *via* air-transportation. To infer the places of origin of those mosquitoes, we genotyped 12 microsatellite loci for which the most comprehensive population genetic reference is available. Although clustering by Bayesian and multivariate methods both suggested all those airport mosquitoes belong to Asia/Pacific population, they were not clustered into a single population. Also, there was variation in mitochondrial *Cox1* haplotypes among mosquitoes collected in different incidents of discovery which indicated the existence of multiple maternal origins. Whereas we conclude there is little evidence to support overwintering of *Ae. aegypti* in the airports in this study, special attention is still desired to prevent the invasion of this prominent arbovirus vector.

## Introduction

*Aedes aegypti*, dengue-yellow fever mosquito, distributes in the most part of the tropical and subtropical regions. This species has strong biting preference to humans and is adapted to urbanized environments that make them an extremely effective vector of numerous arthropod-borne viral diseases including dengue, yellow fever virus and zika virus. The current global distribution of this species is considered a result of intercontinental movement of the mosquitoes along with trades and traffic by human. Recent population genetic studies indicate existence of two major genetic clusters of *Ae. aegypti* world-wide (Brown et al. 2011, Gloria-Soria et al. 2014). The African cluster which distribute exclusively in Africa is believed to be representing the ancestral population of this species. On the other hand, *Ae. aegypti* population distributing in all regions outside of Africa as well as some parts of Africa is a monophyletic population lineage. Mosquitoes in this “out-of-Africa” cluster is more domesticated and well adapted to human inhabitation than mosquitoes in the “Africa” cluster. The out-of-Africa cluster may has been derived from Africa to the other part of the world probably around the 16^th^ century along with transatlantic traffic (Powell et al. 2018).

Modern global transportation may accelerate spread of such important insect pests across continents. For disease transmitting mosquitoes, aircrafts is one of the most important pathway for its daily volume and speed (Gratz et al. 2000, Ibañez-Justicia et al. 2017). In Japan, the number of international scheduled flights increases constantly in these years (Japan Ministry of Land, Infrastructure, Transport and Tourism https://www.mlit.go.jp/koku/koku_fr19_000005.html) partly due to the rapid growth of low-cost carrier business. Thus, reinforcement of surveillance system for the exotic mosquitoes within airports is highly demanded. In 2012, *Ae. aegypti* larvae were discovered in a single oviposition trap placed in a passenger terminal of Narita International Airport (NRT), Chiba, Japan, which was the first detection of this mosquito species in a building of international airport, Japan (Sukehiro et al. 2013). Insecticide was sprayed around the area soon after the discovery, and the following intensive survey did not detect more *Ae. aegypti* in that season (Sukehiro et al. 2013). After the incident in 2012, however, the mosquitoes were sporadically trapped again by oviposition traps (ovitraps) installed in NRT in August and September 2013, September 2014 and June, September and November 2015 (Table 1). Also, in September 2013, single *Ae. aegypti* adult was captured in another airport, Tokyo International Airport (aka Haneda Airport: HND), which locates approximately 60 km south-west away from NRT (Fig. 1). Although the continuous discoveries of *Ae. aegypti* in airports would represent repeated introductions from one or several foreign regions, we also concerned a possibility that there was a source population of *Ae. aegypti* which is overwintering in the airport buildings especially in the cases of NRT where multiple incidents of discovery were recorded.

**Table 1.** Description for incidents in which *Ae. aegypti* were captured in the two international airports, Japan, during 2012—2015.

| ID | Location | Period | Description* | N | Cox1<br>Access. No. |
| --- | --- | --- | --- | --- | --- |
| NRT12 | NRT | 2012 Aug | Pupae and larvae were discovered in single ovitrap. Intensive survey after the discovery in area 400 m around did not detect additional mosquitoes (Sukehiro et al., 2013). | 4 | LC482631 |
| HND13 | HND | 2013 Sep | Single adult male was captured in CDC light trap placed in cargo terminal. Intensive survey after the discovery in area 400 m around did not detect additional mosquitoes | 1 | LC482636 |
| NRT13_Aug | NRT | 2013 Aug | Pupae and larvae were discovered in single ovitrap set in airplane arrival terminal which located 1.5 km depart from NRT13_Sep_Aug discovered spot. Intensive survey after the discovery in 400 m area around the spot did not detect additional mosquitoes. | 8 | LC482632 |
| NRT13_Sep | NRT | 2013 Sep | Pupae and larvae were discovered in single ovitrap set in cargo terminal which located 1.5 km depart from NRT13_Sep_Aug discovered spot. Intensive survey after the discovery in 400 m area around the spot did not detect additional mosquitoes. | 10 | LC482633 |
| NRT14 | NRT | 2014 Sep | Larvae were discovered in single ovitrap. Intensive survey after the discovery in 400 m area around the spot did not detect additional mosquito. | 7 | LC482630 |
| NRT15_Jun | NRT | 2014 Jun | Larvae were discovered in single ovitrap. Intensive survey after the discovery in 400 m area around the spot did not detect additional mosquitoes. | 4 | LC482634 |
| NRT15_Sep | NRT | 2014 Sep | Larvae were discovered in single ovitrap. Intensive survey after the discovery in 400 m area around the spot did not detect additional mosquitoes. | 2 | LC482629 |
| NRT15_Nov | NRT | 2014 Nov | Larvae were discovered in single ovitrap. Intensive survey after the discovery in 400 m area around the spot did not detect additional mosquitoes. | 4 | LC482635 |

\*From Vector Surveillance Reports by Quarantine Information Office, Ministry of Health, Labor and Welfare Japan: https://www.forth.go.jp/ihr/fragment2/index.html)

**Fig. 1.**
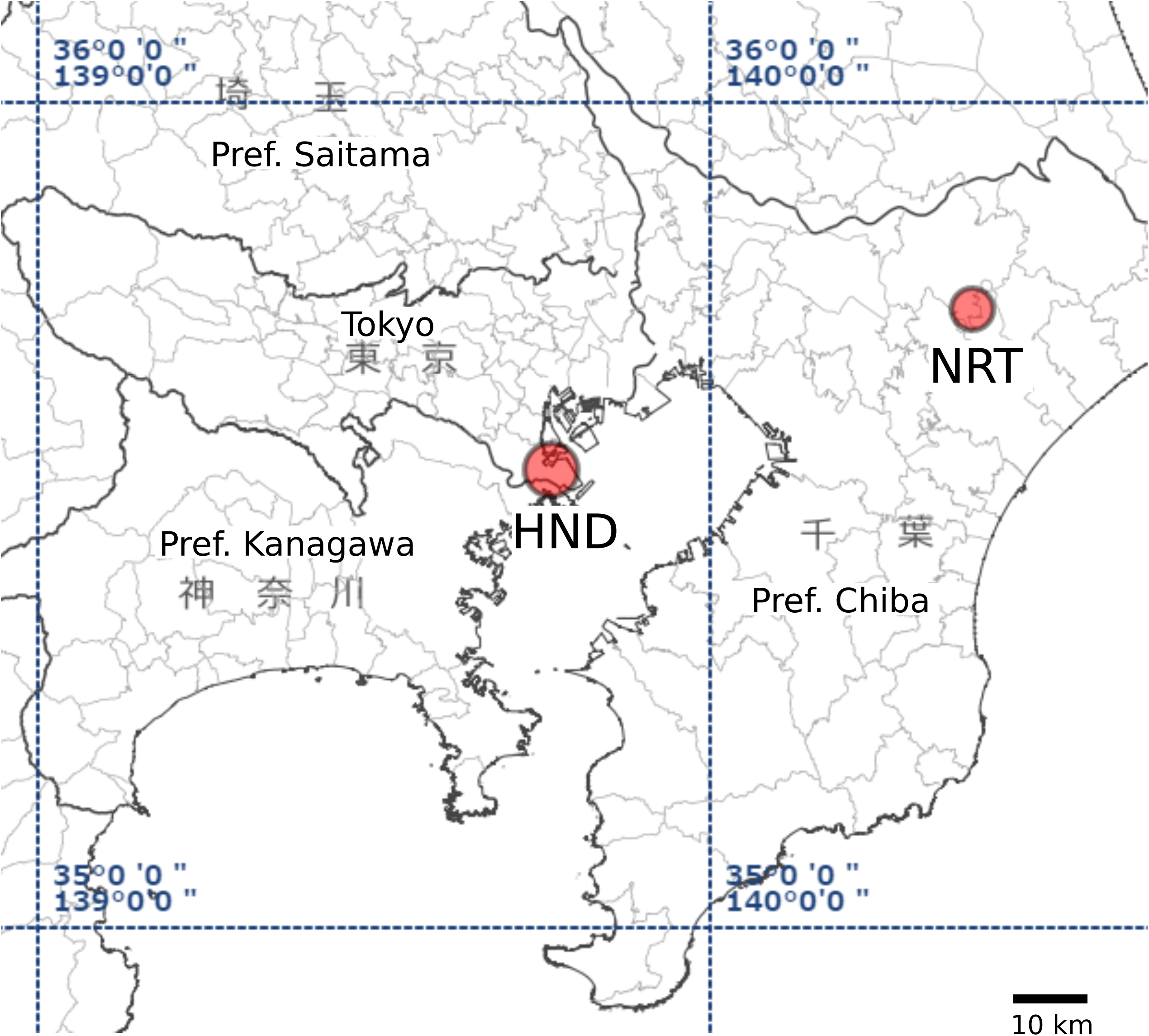
Locations of Narita-airport (NRT) and Haneda airport (HND) on map including Tokyo and peripheral cities. The map was reproduced from Geospatial Information Authority of Japan website (https://www.gsi.go.jp).

Preceding studies using genotype data at 12 microsatellite loci have revealed the hierarchical structure of the world-wide *Ae. aegypti* population (Brown et al. 2011, Gloria-Soria et al. 2016). According to the result of those studies, the world-wide *Ae. aegypti* population is divided into Africa and out-of-Africa clusters, as mentioned above. The out-of-Africa cluster is further divided into two New-world clusters and an Asia/Pacific cluster (Gloria-Soria et al. 2016). We considered such a hierarchical structure and existing comprehensive microsatellite genotype table for this species (Gloria-Soria et al. 2016) allow us to narrow down the origin of the *Ae. aegypti* discovered in airports. In this study, we analyzed genotypes of the 12 microsatellite loci and the sequence of mitochondrial cytochrome oxidase 1 (*Cox1*) gene haplotypes in those mosquito samples captured in airport buildings during 2012—2015.

## Material and methods

### Mosquitoes

Routine mosquito surveillances during 2012—2015 conducted by the airport quarantine station stuffs discovered *Ae. aegypti* larvae and adults in two international airports serving to the Greater Tokyo Area (Fig. 1) (Vector Surveillance Reports by Quarantine Information Office, Ministry of Health, Labor and Welfare Japan: https://www.forth.go.jp/ihr/fragment2/index.html). Among those incidents listed in Table 1, the incident in NRT, 2012 has already detailed in Sukehiro et al. (2013). Those mosquitoes were identified as *Ae. aegypti* morphologically. Some of the larvae were kept in laboratory and grown to adults before being provided to us. For some mosquitoes, we obtained only one or few legs from the quarantine office after the rest of bodies was subjected to flaviviruses and Chikungunya virus detection by RT-PCR.

### DNA extraction

Modified alkaline lysis method (Rudbeck and Dissing 1998) was used to prepare PCR template from one to three legs from single adult mosquito. In our modified alkaline lysis method, legs were homogenized in 10 μl of NaOH solution (0.2 M) in each well of 8-stripped PCR tubes by shaking with a zirconia bead (2 mm in diameter, Nikkato, Japan) in TissuLyser II (Qiagen) for 30 s at 30 Hz. The homogenate was incubated for 10 min on 75 °C, then neutralized by adding 10 μl of neutralization buffer (360 mM tris-HCl, 10 mM EDTA, pH 8.0) and 90 μl of Milli-Q water.

### Genotyping microsatellite loci

Twelve microsatellite loci developed in Brown et al. (2011) and Slotman et al. (2007) were amplified by PCR with labeled M13 primers as described in Brown et al. (2011). Primers and fluorescent dye combinations we used are described in Table S1. A PCR mixture contained 1 ul of template DNA, 1× Type-it Multiplex PCR Master Mix (Qiagen), 0.2 μM of each locus specific reverse primers, 0.02 μM of each locus specific forward primers and 0.2 μM of fluorescent labeled M13 primers, and the PCR condition was 95 °C for 2 min, 40 cycles of 98 °C for 5 s, 55 °C for 90 s and 72 °C for 20 s, then final extension on 72 °C for 1 min. The resulted PCR fragments were electrophoresed with GeneScan 500 LIZ size standard (Applied Biosystems, ABI) in ABI3130 (ABI) for fragment analysis. Allele sizes were scored in Peak Scanner Software v1.0 (Thermo Fisher Scientific). Five DNA samples previously analyzed by Brown et al. (2011) was also genotyped in same manner to calibrate the consistency of allele-call between the different laboratories.

### Population clustering and assignment

The genotype data of airport populations were merged with the reference individual genotype table (VBP0000138 in Population Biology Project of VectorBase.org) (excluding the *Ae. mascarensis* and *Ae. queenslandensis* data) and formatted for analysis by STRUCTURE 2.3.4 (Pritchard et al. 2000, Falush et al. 2003). Each run was conducted with 200,000 burn-in followed by 500,000 sampling, without using prior information of collection locations and with allele frequency correlated model for ten independent runs as replication. The best K value was determined according to the Evanno’s criteria (Evanno et al. 2005). The replications at the best K were averaged by CLUMPP (Jakobsson and Rosenberg 2007), and then visualized by DISTRUCT (Rosenberg 2003) using CLUMPAK server (Kopelman et al. 2015).

Discrimination analysis of principle component (DAPC) was conducted for microsatellite genotype data using adegenet v2.0.1 package (Jombart et al. 2010) in R v3.3.2. Countries of origins (except Hawaii/USA, which were treated as separated regions) were used for predefinition of populations for reference genotype panels. For airport samples collected in Japan, samples collected in each different incident are treated as distinct predefined populations.

Population assignment analyses were conducted in GeneClass2 (Piry et al. 2004). First, the reference genotypes were divided into African and out-of-Africa groups. In self-assignment test, as setting cutoff threshold probability to 0.8, GeneClass2 assigned 95.4% African genotypes (42/918) and 98.0% out-of-Africa genotypes (2661/2714) back to each original group. Misassignment (assigning to wrong cluster with probability >0.8) rates, on the other hand, was 2.9 and 1% for African and out-of-Africa genotypes, respectively. Then, the out-of-Africa reference genotypes were divided into New-World and Asia/Pacific groups. In self-assignment test among these groups, GeneClass2 assigned 88.2% (1740/1972) New-world genotypes and 88.1% (654/742) Asia/Pacific genotypes were properly assigned back to each original group. Misassignments, on the other hand, occurred in 8.3 and 7.3 % for New-world and Asia/Pacific genotypes, respectively.

### Sequencing cytochrome oxidase I gene (*Cox1*) in mitochondrial DNA

The fragments of the *Cox1* genes were amplified individually using primers COI-FOR 5□-GTAATTGTAACAGCTCATGCA-3’/COI-REV 5□-AATGATCATAGAAGGGCTGGAC-3’ (Paupy et al. 2012a). The 10 μl reaction mixes contained1 μl of 10× reaction buffer (Qiagen), 0.8 μl dNTP, 20 pmol of each primer and 1 U of Taq polymerase (Qiagen, USA) and 1 µl of the DNA template. PCR was performed under the following conditions: 94°C for 3 min and 35 cycles of 94°C for 15 s, 55°C for 30 s, 72°C for 30 s; and 72°C for 10 min. The amplified PCR product were cleaned using ExoSAP-IT (USB Corporation, Cleveland, OH, USA) and sequenced in 3730 DNA Analyzer (Applied Biosystems) using BigDye Terminator v 1.1 Cycle Sequencing Kit (Applied Biosystems). The *Cox1* haplotype of NRT13_Sep was queried in BLASTN (Altschul et al. 1990) (2019/05/09) search at NCBI Nucleotide collection (nt/rt) database restricted in *Ae. aegypti*, and hits with more than 95% query coverage were aligned using MUSCLE (Edgar 2004). Sequences retrieved were AF380835, AF390098 and AY056597 (Morlais and Severson 2002); AF425846 (Mitchell et al. n.d.); AY432106 and AY432648 (Bartholomay et al. 2004); EU352212; HQ688292-688298 (Fort et al. 2012), JQ926676-926684, JQ926686-926690, JQ926692-926696, JQ926698-926700 and JQ926702, JQ926704 (Paupy et al. 2012b); KF909122 (Seixas et al. 2013), KM203140-203248 (Jaimes-Dueñez et al. 2015); KT313642, KT313645, KT313648, KT313650-313653 (Calvez et al. 2016); KT339661 and KT339679-339683 (Vadivalagan et al. 2016), KU186990; KX171382-171394. Haplotype network was drawn for the alignment using the *pegas* package (v0.11) in R (Paradis 2010).

## Result

### Population genetic analysis

The 12 microsatellite loci were genotyped for 40 *Ae. aegypti* samples collected in two international airports serving to Greater Tokyo Area during 2012—2015. As already confirmed in preceding study (Brown et al. 2011, Gloria-Soria et al. 2016), STRUCTURE separated the whole individual genotypes (samples in this study + references) into Africa and out-of-Africa genetic clusters at the best K-value=2 with some level of admixture in Kenyan and Argentina (Fig. 2A). All samples collected in the airports belonged to the out-of-Africa cluster. The genotypes from out-of-Africa countries plus the airport samples were further separated into two New-World clusters and one Asia/Pacific cluster at the best K-value=3 (Fig. 2B) as expected from the result of the preceding study (Gloria-Soria et al. 2016). Genotypes of the airport samples showed preferences to the Asia/Pacific cluster. The Asia/Pacific group plus the airport samples were separated at the best K-value=5 (Fig. 2C). The result showed weak population structure according to geographic locations/countries. The airport population in Japan showed affinities to several different clusters. Especially, NRT12 and NTR15_Jun, NRT13, NRT14 and the sole HND13 individual were clustered into Australia, Vietnam-Hanoi, Thailand and Middle-East/Sri Lanka clusters, respectively, with relatively high posterior probabilities (Fig. 2C). DAPC analysis also supported a membership of the airport samples to the out-of-Africa group (Fig. 3A). Clustering genotypes excluding Africa data marginally separated New-World and Asia/Pacific genotypes with substantial overlap. All airport samples were contained in a range of Asia/Pacific cluster (Fig 3B) but were not completely distinct from the New-World cluster. No more fine clustering was obtained from DAPC analysis within Asia/Pacific group (Fig. 3C).

**Fig. 2.**
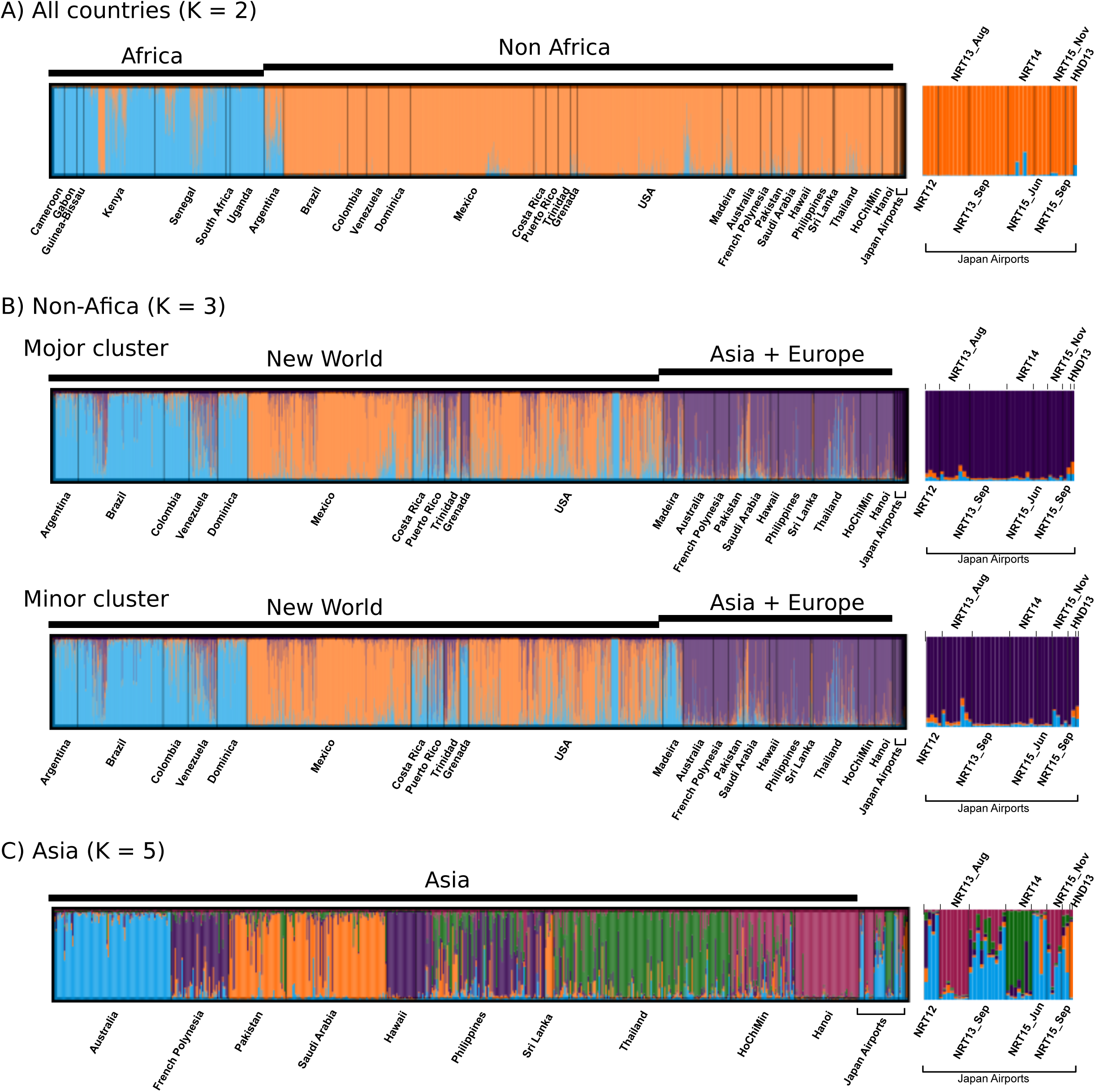
Bayesian clustering by STRUCTURE. Result of multiple STRUCURE runs were averaged by CLUMPP. Only results for the best K-values in Evanno’s method are shown. Magnified views for the airport samples are shown at the right end of each figure. (A) Clusters of all *Ae. aegypti* genotypes + airport samples in Japan at the best k-value 2. (B) Clusters of out-of-African genotypes + airport samples in Japan at the best k-value 3. There were two distinct clustering results. (C) Clusters of Asia/Pacific genotypes + airport samples in Japan at the best k-value 5.

**Fig. 3.**
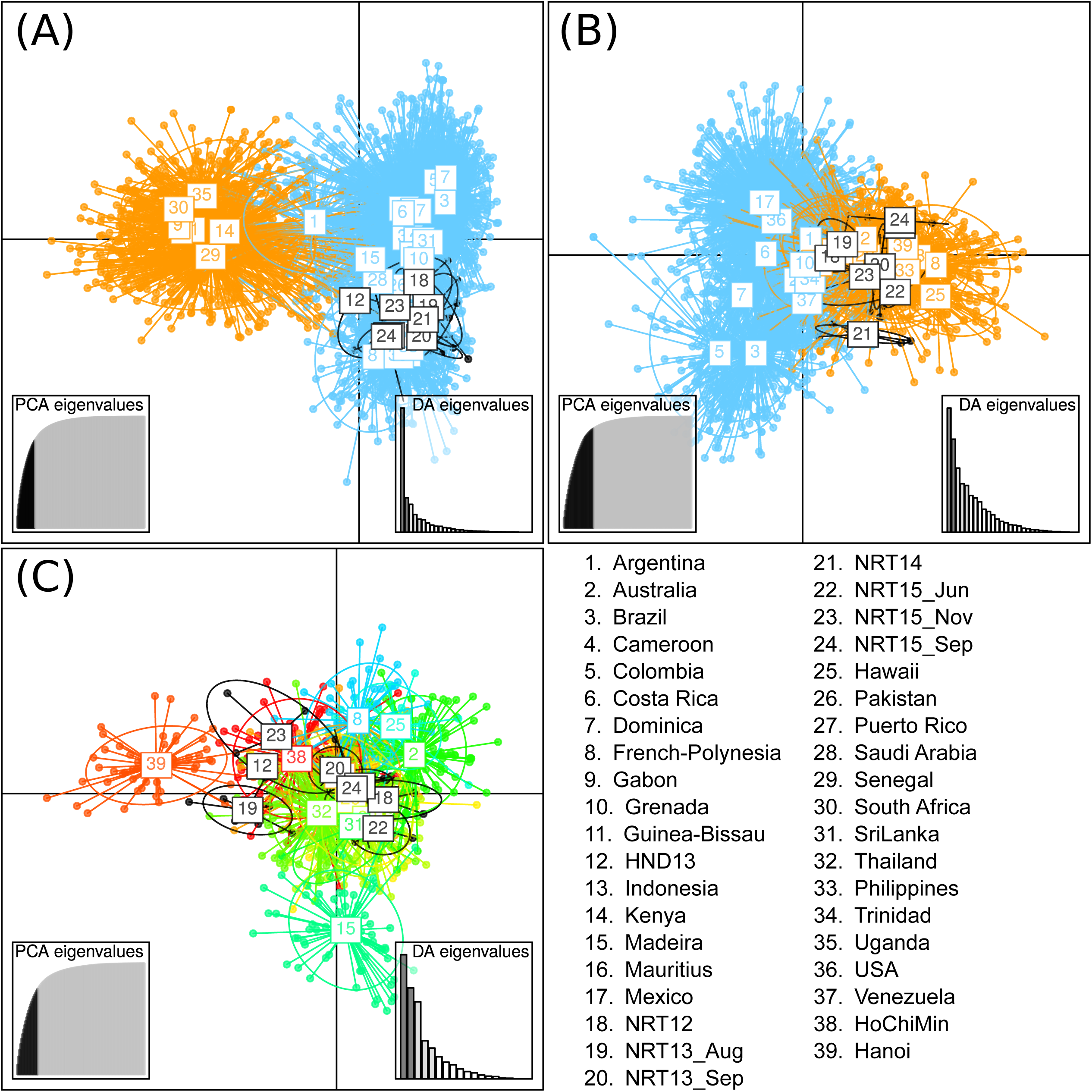
DAPC analysis by Adegenet. The results of Discriminative Analysis of Principle Components (DAPC) are shown. X- and Y- axes indicate 1^st^ and 2^nd^ principal component of DAPC, respectively. Right and left insects show scree plots of PCA and DA eigenvalues, respectively. Labels and individual points for airport samples in Japan are drown in black. First, we conducted clustering using whole genotype data along with airport samples in Japan (A). Next, out-of-African genotypes along with airport samples in Japan were clustered. (B). Finally, Asia/Pacific genotypes along with airport samples in Japan were clustered (C).

The result of STRUCTURE and DAPC was cross validated by assigning the airport samples to *a priori* defined genetic group by GeneClass2. All airport samples, except one in NTR14 assigned to the out-of-Africa group for “Africa or out-of-Africa” selection panel. When using the “New-World or Asia/Pacific” selection panel, most individual genotypes in NRT samples showed Asia/Pacific origin. The sole individual from HND13, on the other hand, was assigned to New-World group (Fig. S1).

### Mitochondrial lineage

Mitochondria *Cox1* gene was sequenced for the 40 *Ae. aegypti* individuals captured in airports. Individuals captured in the same incident had each identical haplotype suggesting the mosquitoes from each incident represent siblings from same female mosquitoes. Fig 4 shows haplotype network graph for the *Cox1* haplotypes of each incident plus other entries retrieved from NCBI Nucleotide collection database. NTR13_Sep, NRT15_Jun and NRT15_Nov mosquitoes shared the same haplotype which was also identical to haplotypes already reported from Asia/Pacific region (Fig. 4). Although other airport samples had each unique haplotype among all airport samples, the haplotypes in HND13 and NRT14 were identical to haplotypes already reported from Asia/Pacific region and both New-World and Asia/Pacific region, respectively.

**Fig. 4.**
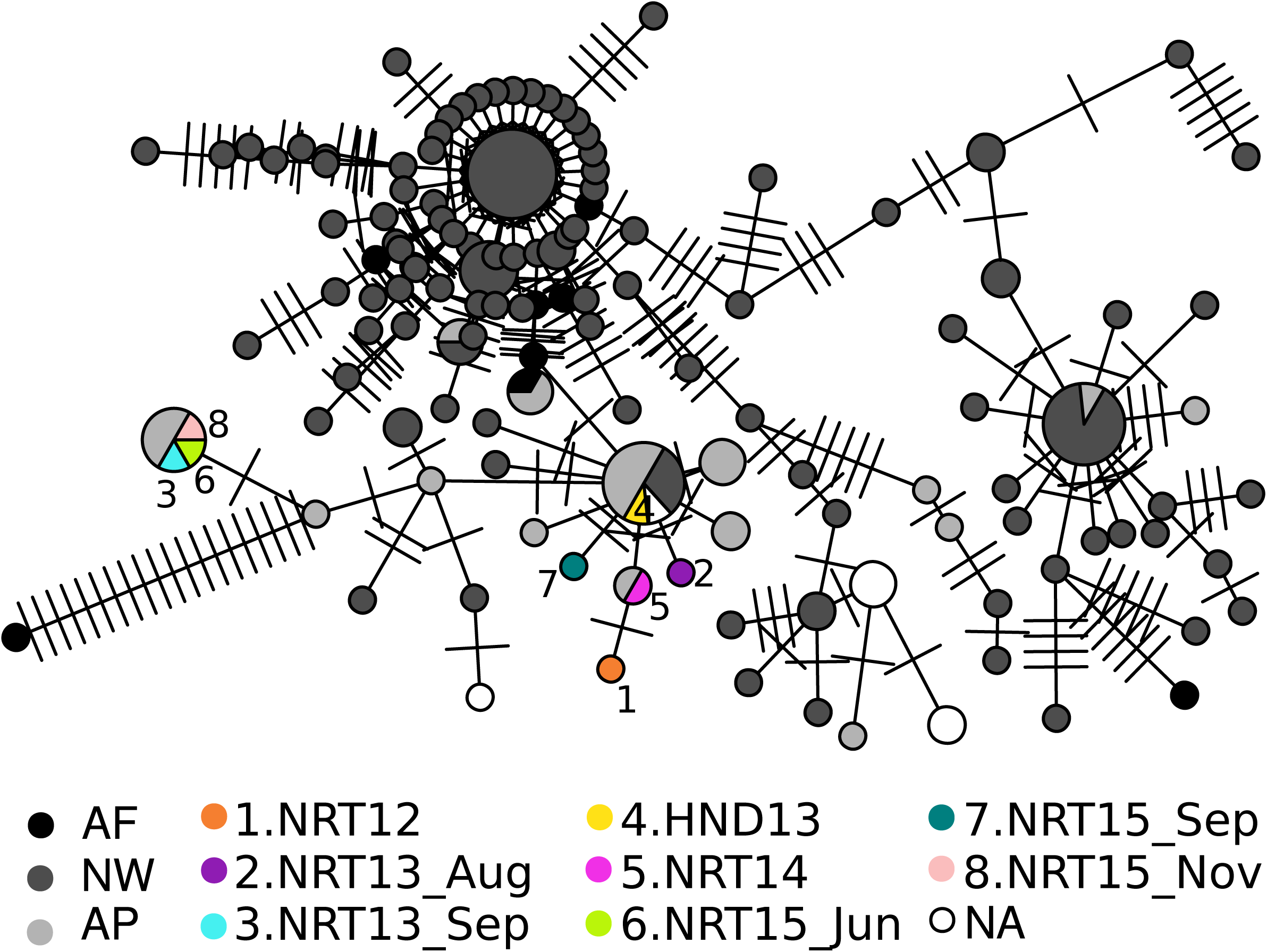
Haplotype network graph for *Cox1* gene. Each node indicates distinct *Cox1* haplotype. Number of ticks on each edge show number of mutations. AF: Africa, NW: New-world, AP: Asia/Pacific, NA: Information unavailable.

## Discussion

### Origin of *Ae. aegypti* captured in Narita and Haneda international airports

We analyzed *Ae. aegypti* sampled in two international airports serving to the Greater Tokyo Area. Both Bayesian (STRUCTURE) and multivariate (DAPC) clustering methods supported all those individuals belong to Asian/Pacific genetic group. Although GeneClass2 assigned most individuals to the Asia/Pacific group by hierarchical clustering approach, one individuals was clustered into Africa cluster in the “Africa or out-of-Africa” selection panel and one NRT15_Nov, and the sole HND13 individual were clustered into New-World cluster in “New-World or Asia/Pacific” selection panel with high probability (>80%) (Fig. S1). During 2012 to 2015, more than half of total passenger planes arriving at Narita Airport originated from Asia/Pacific region every year, while direct flights originating from Africa or South America (most likely source in the New World) accounted for less than 0.3% of the total flights (Sukehiro et al. 2016). Considering relatively high misassignment rate in GeneClass2 test (see Materials and Methods) and the high traffic volume from Asia/Pacific regions to Japan, we, at the moment, assume the origins of all airport samples are somewhere in Asia/Pacific region. Although STRUCTURE analysis clustered some individuals into more specific clusters (Fig. 2C) with relatively high posterior probability, these results should be kept in speculative because number of Asian and Pacific countries represented in the reference panel are still limited. To obtain more confidence and resolution to assign individuals into narrower local populations (i.e. country level), further expansion of reference panel to include more world-wide populations and utilization of richer genetic information such as genome-wide SNPs (Rasic et al. 2014, Evans et al. 2015, Schmidt et al. 2019) will be required.

### Are *Ae. aegypti* reproduce stably in airport?

The mosquitoes collected in same incident had all identical mitochondrial haplotypes with each other. This suggests individuals collected in same incident represented siblings from single female. On the other hand, mosquitoes collected in different incidents could had different mitochondrial haplotypes indicating that there were multiple different maternal lineages for *Ae. aegypti* collected in airports during 2012—2015. Furthermore, STRUCTURE analysis did not assign all airport individuals into single cluster within Asia/Pacific group. Considering the facts that the discovery were occasional and intensive surveys following each discovery did not find additional *Ae. aegypti*, there is so far little evidence to support establishment of stable *Ae. aegypti* population in airport.

While most region in Japan are not suitable for *Ae. aegypti* inhabitation, this species are once established overwintering population in temperate zone in Japan within limited period after the World War II (1944—1952) (Tanaka et al. 1979). Overwintering of *Ae. aegypti* was also suspected in Washington, DC during 2011—2014, where the mosquito may be utilizing the subterranean habitat (Lima et al. 2016). In 2014, local infection of dengue occurred in Tokyo (Kutsuna et al. 2015) for the first time in those 70-years, though the vector mosquito was *Ae. albopictus*. Nevertheless, continuous introduction of both vectors and pathogens pose an undesirable risk that would change epidemiological situation in this country. Thus, further intensive surveillance and preventive measure for exotic mosquito in airport are desired.

## Acknowledgement

This research was partially supported by the Research Program on Emerging and Re-emerging Infectious Diseases from Japan Agency for Medical Research and development (AMED) and by Japan Initiative for Global Research Network on Infectious Diseases (J-GRID) from Ministry of Education, Culture, Sports, Science and Technology in Japan and AMED. We acknowledge to stuffs in the Quarantine Stations in Narita and Haneda airports for kindly providing insect samples. Also, we thank to Dr Jeff Powell and Dr. Gloria-Soria in Yale university for providing reference *A. aegypti* genomic DNAs for electrophoresis calibration.

