## Supplemental Figure S1 for "Genetic analysis of *Aedes aegypti* captured in two international airports serving to the Greater Tokyo Area during 2012—2015"

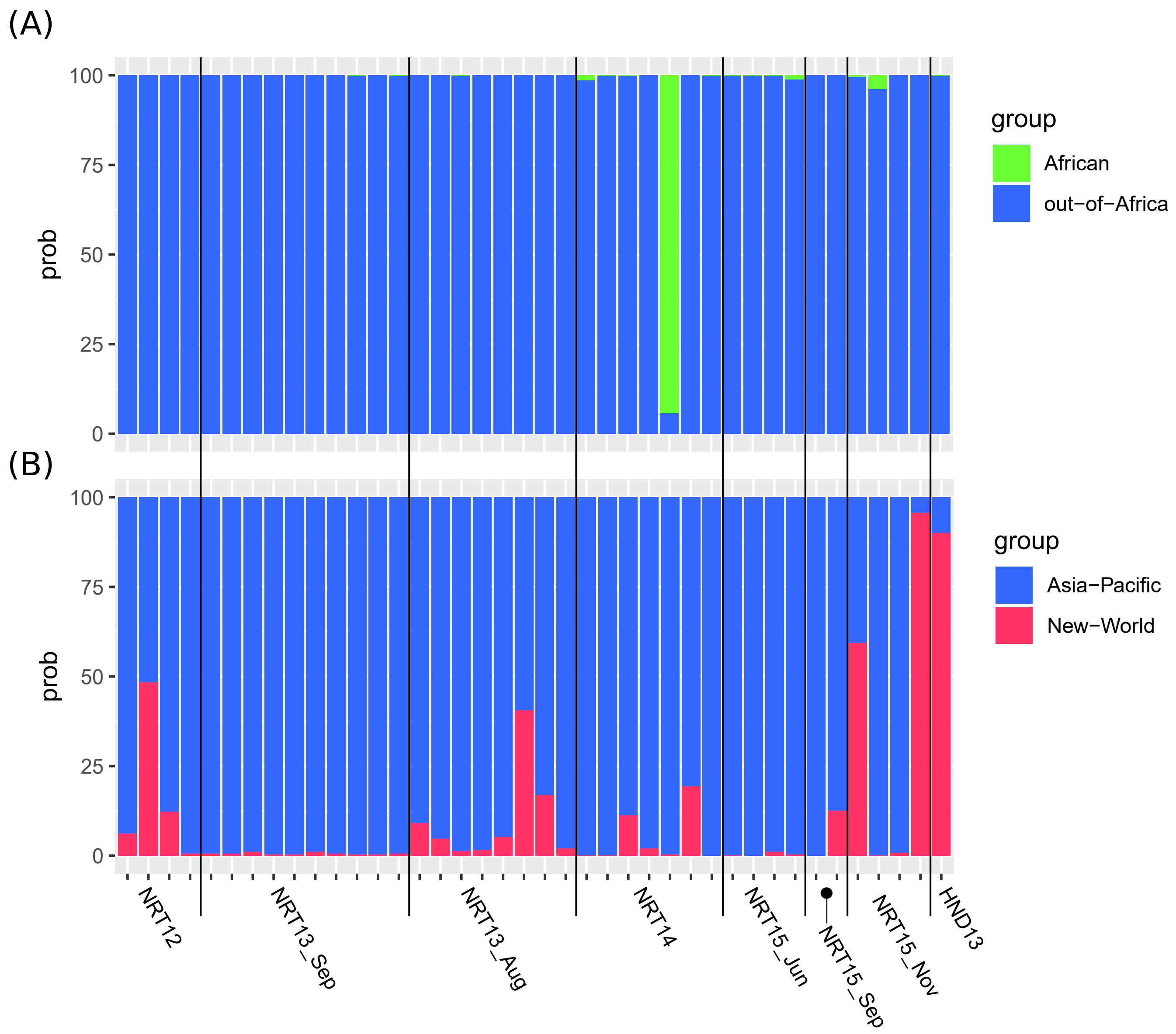


### Fig. S1 Population assignment experiment

GeneClass2 was used to assign genotypes of airport samples to predefined population groups. Each bar indicates posterior probability of assignment to each population group of each individual. (A) Predefined population groups were Africa/out-of Africa. (A) Predefined population groups were New World/Asia-Pacific.
